# *Yellow* shapes mosquito pigmentation and vector competence to *Plasmodium berghei*

**DOI:** 10.1101/2025.09.24.678207

**Authors:** Dennis Klug, Amandine Gautier, Eric Marois, Stéphanie Blandin

## Abstract

The insect-specific *yellow* gene, first described over a century ago, remains enigmatic despite its conserved role in pigmentation. In the malaria vector mosquito *Anopheles coluzzii*, *yellow* is strongly expressed during juvenile stages and in adult ovaries, mirroring pigmentation patterns that likely enhance desiccation resistance, camouflage, and reproductive fitness. To dissect its function, we generated a complete knockout allele *yellow(-)KI* by replacing the first exon with an EGFP reporter, and introgressed it into multiple genetic backgrounds. Loss of *yellow* abolished black cuticle pigmentation and delayed larval development but left adult lifespan unaffected. Developmental delay likely impaired mating success in competition with wild-type mosquitoes, hastening the decline of the X-linked *yellow(-)KI* allele. Notably, *yellow(-)KI* females showed a consistent twofold increase in susceptibility to the rodent malaria parasite *Plasmodium berghei*, independent of the complement factor TEP1 and melanization, whereas infection with *P. falciparum* was unchanged. Antibiotic treatment reversed the heightened *P. berghei* susceptibility, implicating altered gut microbiota as a mediator. Our findings reveal that *yellow* not only shapes pigmentation but also indirectly modulates vector competence through microbiota-dependent mechanisms, underscoring the complex interplay between cuticle biochemistry, microbial ecology, and pathogen transmission.

## Introduction

Defects in cuticle or eye pigmentation have long served as powerful genetic markers in *Drosophila melanogaster*, providing visually striking, easily scorable phenotypes that facilitated some of the earliest genetic mapping efforts. Mutations in pigmentation genes were among the first identified in classical mutagenesis screens, as the resulting phenotypes were easy to detect and generally viable under laboratory conditions. The first null allele of *yellow* was described as early as 1911 by Edith Wallace (Lindsley D.L., 1968). In *yellow*-deficient flies, black pigment is absent from the entire adult cuticle as well as from larval mouthparts and denticles (Geyer and Corces, 1987; Wittkopp, True and Carroll, 2002; Hinaux *et al*., 2018). Comparable phenotypes have been reported in *Bombyx mori*, *Tribolium castaneum*, *Apis mellifera*, *Rhodnius prolixus* and other insects, following either germline transgenesis or RNAi-mediated knockdown of *yellow* family genes (Futahashi *et al*., 2008; Noh *et al*., 2015; Nie *et al*., 2021; Berni *et al*., 2022). In *D. melanogaster*, *yellow* is strongly expressed from ∼46 h after pupation in wing discs, thorax, and abdomen, tissues that later develop intense pigmentation, and in specific neuronal populations throughout the life cycle (Hinaux *et al*., 2018). This dual expression pattern underlies phenotypes that go beyond visible pigmentation. While reduced male mating success in *yellow* mutants was initially attributed to behavioural changes linked to neuronal expression (Drapeau, Cyran, *et al*., 2006), recent work demonstrated that decreased pigmentation weakens the stiffness of male sex combs, impairing their ability to grasp females during mating (Massey *et al*., 2019). *Yellow* belongs to a rapidly evolving insect-specific protein family, with up to 14 members in *Anopheles gambiae* and *A. mellifera*, but only eight in *D. pseudoobscura*. Such variation suggests lineage-specific diversification or loss under selective pressure (Drapeau, Albert, *et al*., 2006). While *yellow*-like proteins have also been identified in bacteria such as *Deinococcus radiodurans* (Maleszka and Kucharski, 2000), phylogenetic analyses confirm that the insect family is distinct. Despite over a century of study, the biochemical role of Yellow remains unresolved (Drapeau, 2003). Cuticle pigmentation derives from tyrosine, which is converted to L-Dopa by tyrosine hydroxylase (TH) and then to dopamine by L-Dopa decarboxylase (DDC) (**Fig. 1**). Dopamine acts as the precursor for multiple pigmentation pathways, producing brown NBAD pigment, black eumelanin, yellow pheomelanin, or colourless sclerotization components. The cysteine peptidase Tan reverts NBAD to dopamine, and loss of *tan* reduces pigmentation, though less severely than *yellow* loss (True *et al*., 2005). *Yellow* is thought to act in the conversion of dopamine quinone to eumelanin, either directly or via dopachrome intermediates. Its sequence similarity to a dopachrome conversion enzyme in *Aedes aegypti* (Johnson, Li and Christensen, 2001) led to the hypothesis that it encodes such an activity (Wittkopp, True and Carroll, 2002). While dopachrome-converting activity has been demonstrated for Yellow-f and Yellow-f2, no biochemical evidence exists for Yellow itself (Han *et al*., 2002). Research in *D. melanogaster* has largely focused on Yellow’s role in pigmentation patterns, mate choice, and speciation. More recently, Yellow has also been linked to cuticle stiffness and eggshell permeability (Farnesi *et al*., 2017; Massey *et al*., 2019; Noh *et al*., 2021). However, whether altered pigment precursor homeostasis in non-cuticular tissues has systemic effects remains poorly understood. This is particularly relevant given that melanization is a key immune defence in insects, targeting parasites such as microfilariae in mosquitoes, parasitic wasps in *Drosophila*, and *Plasmodium* ookinetes during their passage through the mosquito midgut epithelium. Here, we demonstrate for the first time that Yellow influences antiparasitic defence in the malaria vector *Anopheles coluzzii*. Knockout of *yellow* increased susceptibility to the rodent malaria parasite *Plasmodium berghei* independently of the complement factor TEP1 and melanization capacity, and this effect could be reversed by antibiotic treatment, implicating the gut microbiota as a mediator. Our findings reveal that a gene classically associated with cuticle pigmentation can exert systemic effects on microbial ecology and vector–parasite interactions, highlighting an unanticipated link between insect pigmentation pathways and pathogen transmission.

**Figure 1.**
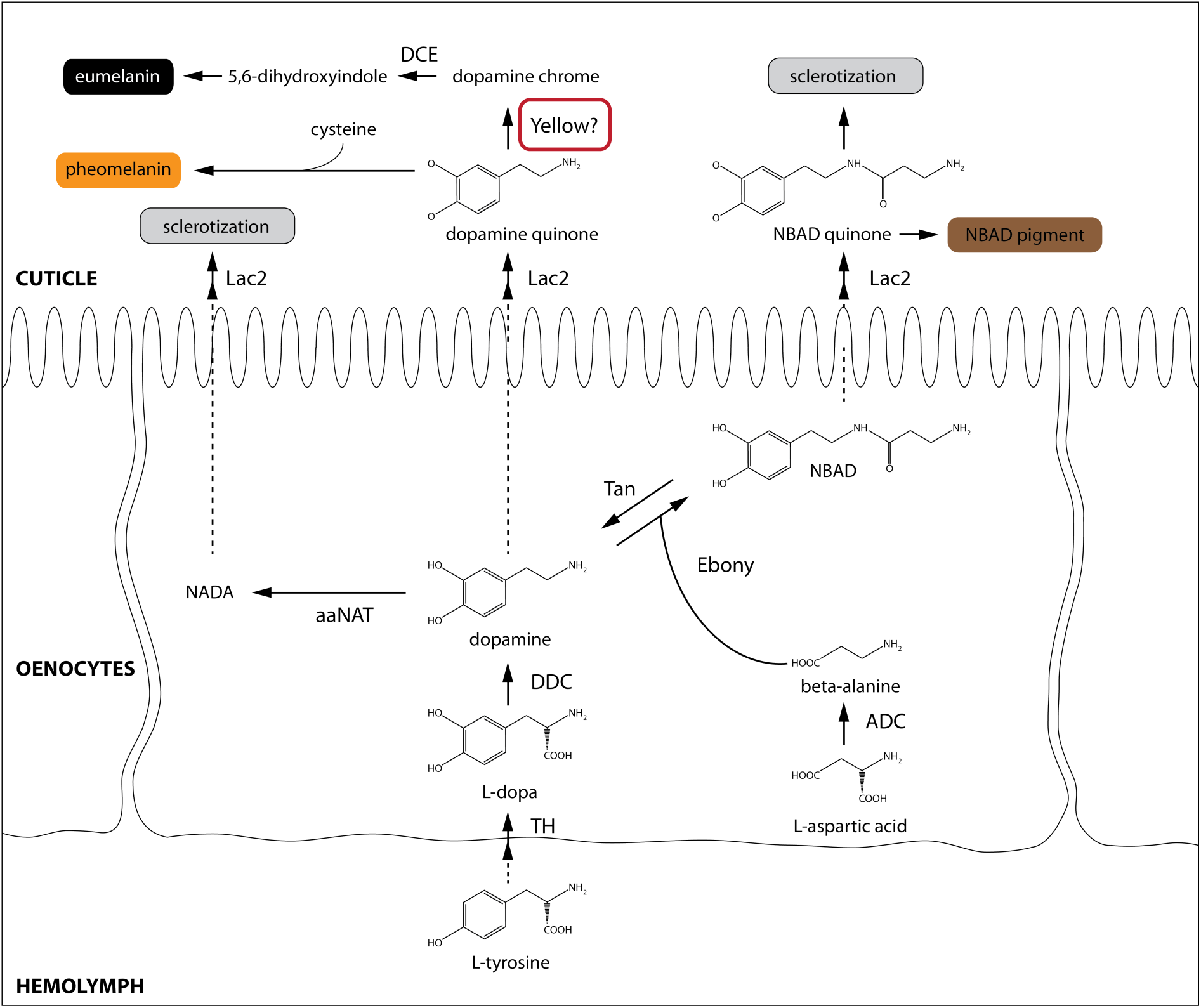
Cuticle pigmentation pathway in insects. Illustration modified from (Pentzold *et al*., 2018). In oenocytes, the amino acid tyrosine is converted by the enzyme tyrosin hydroxylase (TH) into L-Dopa and further decarboxylated by the L-Dopa decarboxylase (DDC) into dopamine. Dopamine is the precursor of multiple pigments. It can be converted into N-acetyl dopamine (NADA) through enzymatic activity of the arylalkylamine-N-acetyltransferase (aaNAT) which is believed to lead to the formation of mostly colorless components of the cuticle required for sclerotization (Osanai-Futahashi *et al*., 2012). Another branch of the pathway leads to the formation of N-ß-alanyl dopamine (NBAD) via conjugation of dopamine to ß-alanine by the ß-alanyl biogenic amid synthetase Ebony (Richardt *et al*., 2003). This enzymatic reaction can be reverted by the cysteine peptidase Tan keeping dopamine and NBAD in equilibrium (Wagner *et al*., 2007). While being secreted from epidermal cells to form the cuticle, NADA, NBAD and dopamine are oxidized by Laccase 2 (Lac2) into dopamine quinone and NBAD quinone, the latter being further processed to NBAD pigment and sclerotizing components of the cuticle. Dopamine quinone is either conjugated to cysteine to form pheomelanin, or further processed into dopachrome and 5,6-dihydroxyindole through activity of a dopachrome conversion enzyme (DCE) to form black eumelanin. Yellow is believed to act during both, or one of the last two steps of the eumelanin formation. L-dopa: L-dihydroxyphenylalanine; NADA: N-acetyl dopamine; NBAD: N-ß-alanyl dopamine; DCE: dopachrome conversion enzyme; Lac2: Laccase 2; aaNAT: arylalkylamine-N-acetyltransferase; DDC: L-dopa decarboxylase; ADC: aspartate decarboxylase; TH: tyrosine hydroxylase. Although Yellow, highlighted by a red square, is believed to act in the last steps of eumelanin synthesis, the exact function of Yellow is still unknown.

## Results

### Phenotypic characterization of *yellow(-)KI* mosquitoes and promoter trap activity

To generate a *yellow* loss of function mutant in the malaria mosquito *A. coluzzii,* we targeted the first exon of *yellow* (AGAP000879) by CRISPR/Cas9 (Dong *et al*., 2018). A repair template was provided to replace the targeted sequence by a fluorescence cassette encoding EGFP under control of the artificial *3xP3* promoter (**Fig. 2A**) (Klug *et al*., 2022). *3xP3* is predominantly expressed in the eyes and ventral nervous chain of larval mosquito stages and enables efficient sorting of transgenic larvae. *3xP3* was flanked by *lox* sites in order to place *egfp* under the control of the endogenous *yellow* promoter upon Cre-mediated recombination. *Yellow(-)KI* mosquitoes were easily identified both through GFP expression and the complete absence of eumelanin as previously described in the fruit fly *D. melanogaster* (Wittkopp, True and Carroll, 2002; Hinaux *et al*., 2018) (**Fig. 2B**). Mutant mosquitoes displayed complete yellow body coloration and appeared more translucent, especially at the abdomen, allowing better imaging through the cuticle (Klug *et al*., 2022). Interestingly, the eggs of *yellow(-)KI* females were completely yellow, in contrast to the black coloration of wild type eggs (**Fig. 2C**). We noticed that, even without removing the *3xP3* promoter, *egfp* was expressed in other tissues than the eyes and nervous chain, where *3xP3* is restricted (**Fig. 2D**, compares EGFP fluorescence in *yellow(-)KI* and *sag(-)KI*). These results suggest that in *yellow(-)KI*, *egfp* expression is also influenced by the directly neighbouring endogenous *yellow* promoter. High levels of EGFP fluorescence were visible in developing *yellow(-)KI* larva, before hatching from eggs (not shown), as well as in all larval stages and in pupae (**Fig. 2D**). EGFP was expressed mainly in the head and at the anterior of each abdominal segment, resulting in a zebra-like appearance. This expression pattern matches that of a *D. melanogaster* line expressing a fluorescently labeled Yellow protein (Hinaux *et al*., 2018). However, in this line, the zebra-like expression pattern first appears in pupae while larvae display little *yellow* expression that is restricted to a subset of neurons, and possibly cells responsible for the development of the larval mouthparts (Hinaux *et al*., 2018). EGFP fluorescence in *yellow(-)KI* larval stages, especially in the abdomen, appears mostly controlled by the *yellow* promoter.

**Figure 2.**
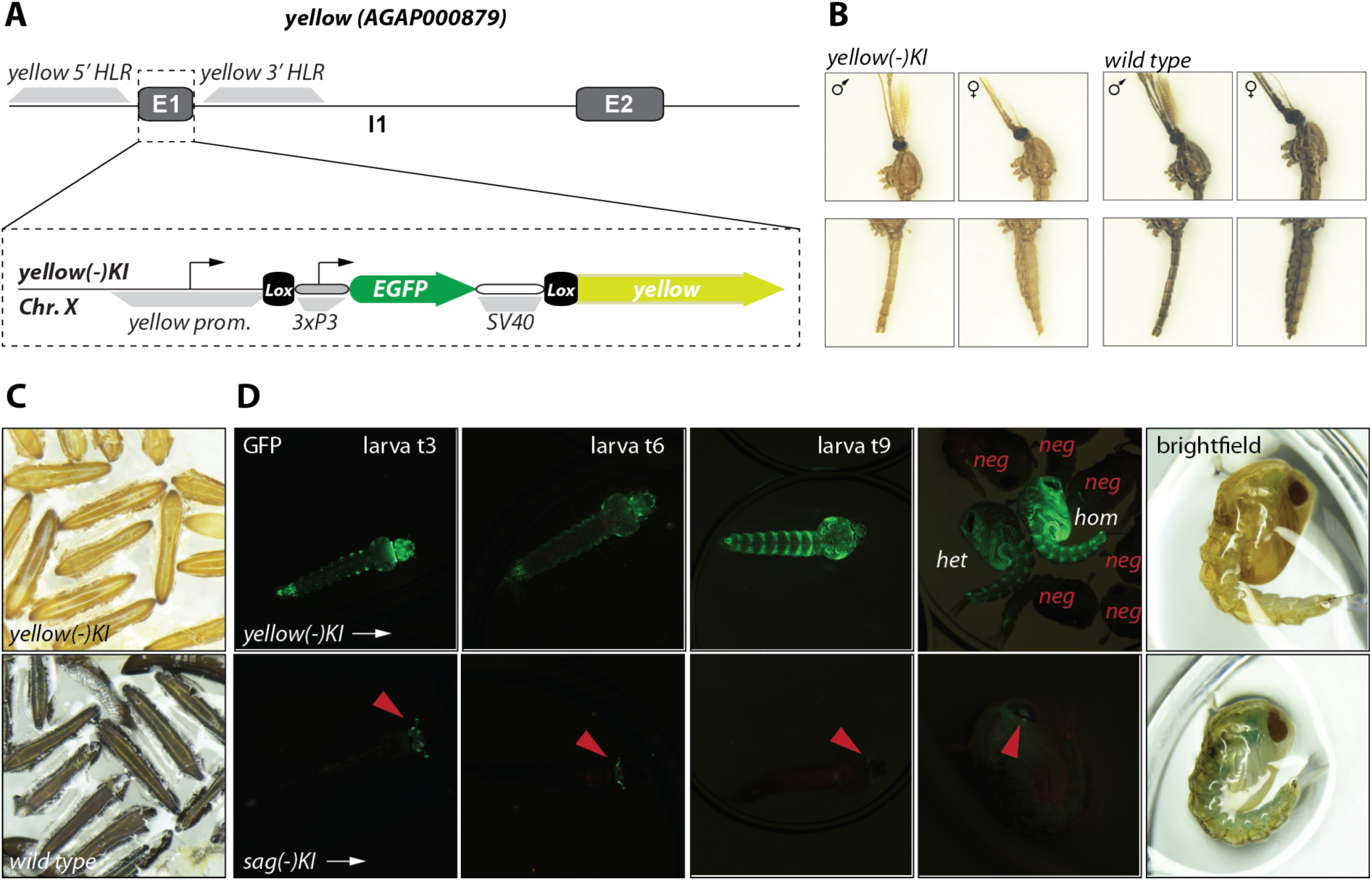
Generation and characterization of the *yellow(-)KI* transgene. **A)** Genomic locus of *AgYellow*. The knockin (KI) transgene is composed of the EGFP cassette flanked by Lox sites. Upon insertion, exon 1, including the start codon, is replaced by the transgene, thus preventing transcription of *yellow*. Transgenic larvae were selected through expression of the fluorescent marker EGFP driven by the *3xP3* promoter. Note that the illustration is not drawn to scale. **B)** Changes in cuticular pigmentation of *yellow(-)KI* females and males compared to wild type. Note the complete absence of eumelanin in *yellow(-)KI* mosquitoes. **C)** *yellow(-)KI* eggs compared to wild type about 24 hours after oviposition (bright field). **D)** EGFP fluorescence in *yellow(-)KI* larvae (top images) at days 3 (d3), 6 (d6) and 9 (d9), and in pupae. EGFP fluorescence and bright field images are shown for the pupal stage. For *yellow(-)KI* a homozygous (hom) and a heterozygous (het) pupae surrounded by negatives are shown. The same developmental stages are shown for the *sag(-)KI* line in which *egfp* is expressed exclusively under the control of the *3xP3* promoter in juvenile stages, when the *Saglin* promoter is transcriptionally silent (Klug *et al*., 2023). Fluorescence signals in this line are concentrated in the head and eyes and become weaker with progressive development (red arrows).

### Fitness costs caused by the absence of Yellow

To distinguish between transgenic and wild type siblings, we made use of flow cytometry based on the larval expression of the introduced fluorescence cassette (Marois *et al*., 2012). Initial analyses of the *yellow(-)KI* line revealed that the X-linked *yellow* gene undergoes dosage compensation, which is reflected by a lower fluorescence in heterozygous females as compared to transgenic males, although both of them carry a single copy of the *yellow(-)KI* cassette (**Fig. 3A,B**). Indeed, by sorting the L1 larvae of each population and rearing them separately to adulthood, we could define the population with the lowest EGFP intensity to be heterozygous females, while males displayed intermediate and homozygous females, the highest fluorescence intensity. To assess potential fitness costs associated with the absence of Yellow, we mixed an equal number of homozygous *yellow(-)KI* and wild type siblings, and we followed the proportion of each genotype over 6 generations (**Fig. 3C**). We observed a sharp decline of the proportions of transgenic larvae (homozygous, hemi- and heterozygous). After six generations, the wild type had almost entirely overtaken the cage population, indicating a high fitness cost for mosquitoes lacking *yellow* expression (**Fig. 3C**). To investigate at which stage of the life cycle the fitness of *yellow(-)KI* carriers was reduced, L1 larvae obtained from wild type and *yellow(-)KI* colonies were mixed in equal ratios and raised until pupation. New pupae were sorted each day according to their EGFP fluorescence and the proportion of wild type / *yellow(-)KI* was calculated (**Fig. 3D**). The first pupae were observed at day 7 after eggs hatched and appeared to be nearly exclusively wild type. The proportion of pupae homozygous for *yellow(-)KI* was always below 50% except on the last collection day (d10) when it reached its maximum (63%). These data indicate that pupation is delayed by ∼2 days in *yellow(-)KI*, and that a larger proportion of *yellow(-)KI* mosquitoes die during juvenile stages as compared to wild type. To identify the stages most sensitive to the lack of Yellow, we daily followed larval and pupal development in different *yellow(-)KI* and WT lines that were raised separately (**Fig. 3E**): *yellow(-)KI*, a double mutant *yellow(-)KI;tan(-)KI* lacking expression of *yellow* and *tan*, a *yellow(-)KI-7b* mutant homozygous for *yellow(-)KI* and the *7b* transgene suppressing the expression of the major complement factor *TEP1* (Pompon and Levashina, 2015), and a *yellow(-)aapp-DsRed* line carrying *yellow(-)KI* together with a fluorescent reporter transgene driving expression of *DsRed* in the outer lobes of female salivary glands (Klug *et al*., 2022). The colonies *yellow(-)KI* and *yellow(-)KI;tan(-)KI* were generated in the *Ngousso* genetic background, while *yellow(-)KI-7b* and *yellow(-)aapp-DsRed* were hybrids of *Ngousso* and *G3L*, another frequently used wild type strain. Both *yellow(-)KI* and *yellow(-)KI;tan(-)KI* showed a strong decrease in the pupation rate, with only 25% larvae reaching the pupal stage for these two lines, vs. 75% for wild type controls. Interestingly, the pupation rate was higher for larvae from the two *yellow(-)KI* lines in the mixed *Ngousso/G3L* background, with *yellow(-)aapp-DsRed* displaying a similar pupation success as wild type (**Fig. 3E**). These results suggest that the fitness costs linked to Yellow loss vary depending on the genetic background and/or its degree of polymorphism. Of note, in both wild type and all *yellow(-)KI* lines, most losses occurred during larval development, followed by the last larval (L4) and the pupal stage (**Fig. 3E, S1**). In *yellow(-)KI* (*Ngousso* backgound) as well as in the two *yellow* lines in the *Ngousso/G3L* background, pupae with moulting defects were also detected (**Fig. S1**). We further measured the life span of females homozygous for *yellow(-)KI* (**Fig. 3F**). For this, both genotypes at L1 larval stage were mixed in equal quantities and raised in synchrony. Four to five days after adult emergence, *yellow(-)KI* and wild type females were separated based on their fluorescence, placed in paper cups and kept under normal breeding conditions. The number of dead mosquitoes was monitored daily until no living mosquito remained. All mosquitoes died within 55 days with no difference in the life span of *yellow(-)KI* and wild type mosquitoes (**Fig. 3F**).

**Figure 3.**
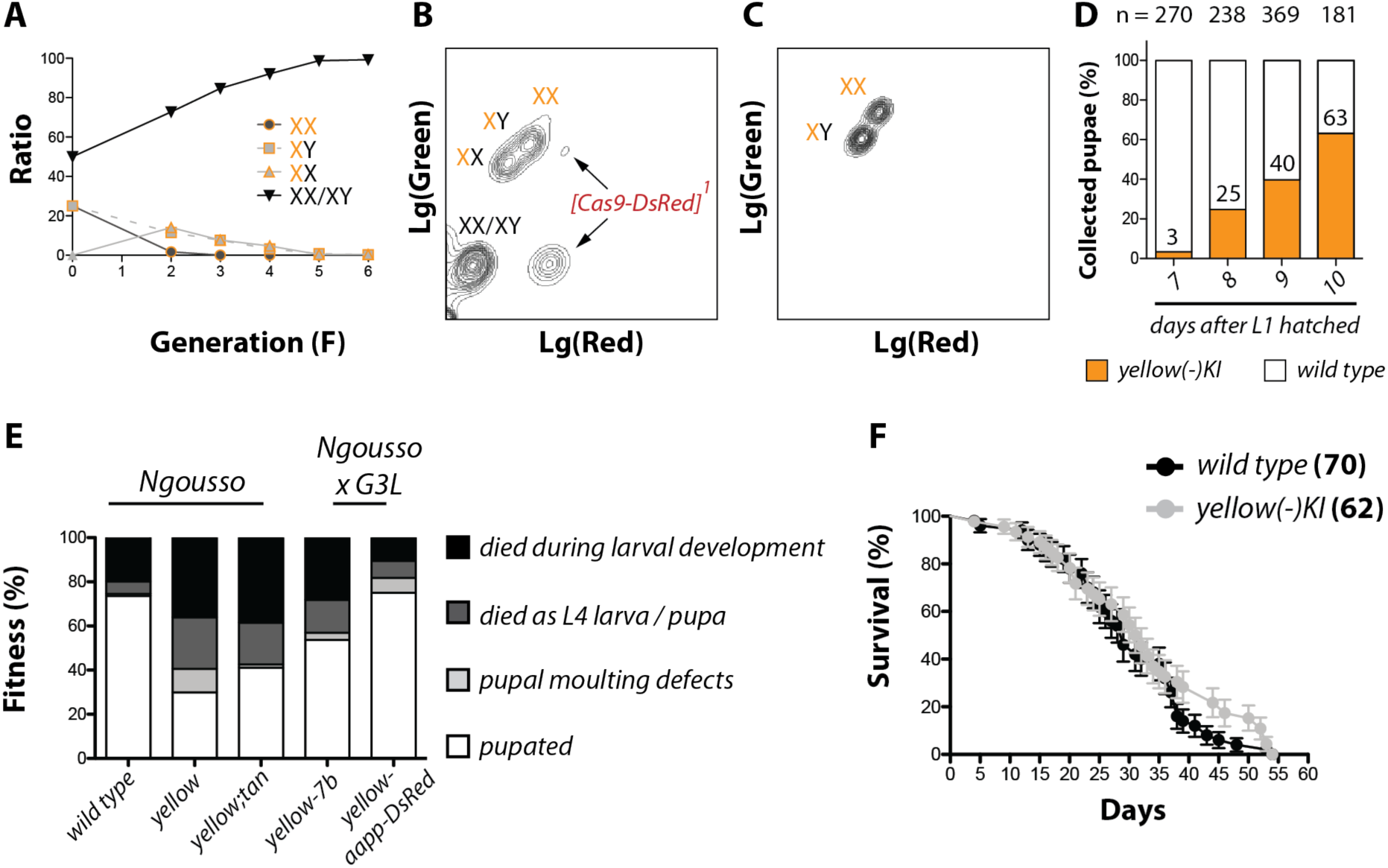
Absence of Yellow delays larval development but does not affect the life span of adult mosquitoes. **A)** The fitness of mosquitoes carrying *yellow(-)KI* was assessed in competition with wild type over six generations. Initially 50 males and 50 females homozygous for *yellow(-)KI* were mixed with 50 male and 50 female transgene-negative siblings. The distribution of different genotypes was monitored over five generations by COPAS analysis of more than 2000 L1 larvae per generation. The F1 generation was not analyzed because of the low number of larvae in this generation. The fact that heterozygous females are less fluorescent than *yellow(-)KI* hemizygous males made it possible to follow these two populations separately. **B)** COPAS plot of L1 larvae of the F3 generation following transgenesis. Larvae carrying the *yellow(-)KI* transgene form three populations; heterozygous females (lowest EGFP intensity), *yellow(-)KI* positive males (medium EGFP intensity) and females homozygous for *yellow(-)KI* (highest EGFP intensity). Larvae carrying no transgene form a population in the bottom left corner. Note that some larvae still inherited the *Cas9:RFP* transgene. As a result additional populations are visible carrying either solely *Cas9:RFP* or *Cas9:RFP* in combination with *yellow(-)KI*. **C)** Flow cytometry of L1 offspring from a mosquito colony homozygous for *yellow(-)KI*. Note that the only remaining populations are homozygous *yellow(-)KI* females and hemizygous males. **D)** Larvae negative or homozygous for the transgene *yellow(-)KI* were mixed in equal numbers and raised until pupation. Pupation started 7 days and ended 10 days after larvae hatched from eggs. Mean percentages of transgenic and wild type pupae from three independent experiments with 400 larvae per experiment and genotype are shown. **E)** Pupation rate of mosquitoes homozygous for *yellow(-)KI* only, or *yellow(-)KI* in combination with homozygous *tan(-)KI*, 7b or *aapp-DsRed* in comparison to wild type mosquitoes. The genetic background of the mosquito colonies is shown above the bars. For each experiment at least 800 larvae per colony were raised synchronously and the number of pupae was counted until day 11 after larvae hatched from eggs. Pupae, dead L4 larvae and pupae, as well as pupae with moulting defects were counted and the percentage in relation to the total population of seeded larvae was calculated. Means of 2 (wild type, *yellow(-)KI* and *yellow(-)-aapp-DsRed*) or 3 (*yellow(-)KI;tan(-)KI* and *yellow(-)KI-7b*) experiments. **F)** Survival of homozygous *yellow(-)KI* females and wild type siblings kept on 10% sucrose solution. The number of dead mosquitoes was monitored daily. Shown is the mean and standard error of the mean from two independent experiments with >30 mosquitoes each. Data were tested for significance with the Log-rank (Mantel-Cox) test. ns: not significant.

### Impact of Yellow on infection with *Plasmodium spp*

To investigate whether *yellow(-)KI* mosquitoes display an altered susceptibility to the rodent malaria parasite *Plasmodium berghei*, wild type (*Ngousso*) and *yellow(-)KI* mosquitoes raised as a mix were infected with high and low doses of *P. berghei* gametocytes (**Fig. 4A,B**). In both cases, *yellow(-)KI* females carried significantly higher oocyst numbers compared to wild type controls (medians in high gametocyte dose: 115 vs 68, and in low dose: 46 vs 27, respectively). To exclude that the increase in susceptibility was linked to the impaired fitness of the *yellow(-)KI* colony (**Fig. 3E**), we repeated infections with high and low doses of *P. berghei* gametocytes using the *yellow(-)aapp-DsRed* colony which displayed a similar pupation rate as wild type (**Fig. 3E**). *Yellow(-)aapp-DsRed* were also more susceptible to *P. berghei* infection than wild types, with more pronounced differences (medians in high gametocyte dose: 121 vs 20, and in low dose: 6 vs 21 in *Yellow(-)aapp-DsRed vs WT*, respectively, **Fig. 4C,D**). In contrast, no significant difference in the parasite load could be detected between *yellow(-)KI* and wild type upon *P. falciparum* infection (**Fig. 4E)**. Albeit not statistically significant, *yellow(-)KI* females even appeared to be less infected than WT (medians: 7 vs 11, respectively), possibly indicating that the absence of Yellow has a negative effect on the infection with human malaria parasites as previously shown for the knockdown of *Yellow-g* in *Anopheles dirus* upon infection with *P. vivax* (Mongkol *et al*., 2021).

**Figure 4.**
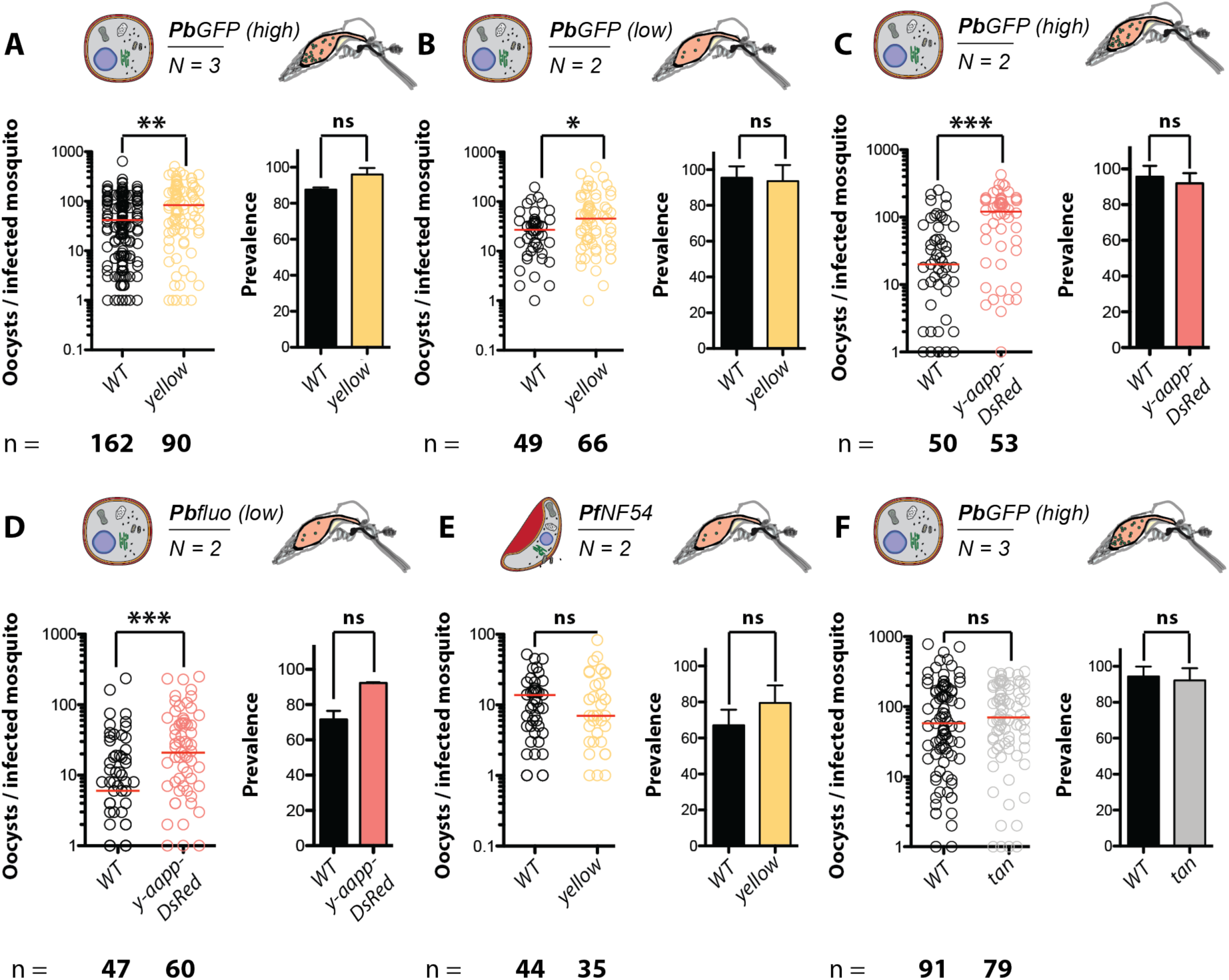
The loss of *yellow* but not of *tan* increases parasite load upon infection with ***Plasmodium berghei* but not with *Plasmodium falciparum*.** For each experiment, WT and sibling control mosquitoes were infected with *P. berghei* (*PbGOMO* or *Pbfluo*) or *P. falciparum* (*PfNF54*). The mosquito line (*yellow(-)KI*, yellow; *tan(-)KI*, gray, *yellow(-)KI + aapp-dsRed*, red; *WT* controls, black) is indicated below the graphs and the *Plasmodium* species and strain, and infection regime (high or low dose) is depicted above each dataset. For each mosquito line and parasite regime, the number of oocysts per infected mosquito is plotted, with red lines indicating medians. The percentage of infected mosquitoes (prevalence) is shown in the bar graph as the mean ± SEM of N replicates. The number of biological replicates (N) is shown below the pictogram while the total number of analysed mosquitoes is shown below each genotype. Statistical significance was tested using Mann Whitney (MW) to compare parasite loads, and Fisher’s exact test (FT) for prevalences. **A)** MW: **p = 0.0015; FT: not significant (ns), p = 0.9206. **B)** MW: *p = 0.0245; FT: ns, p = 0.8374. **C)** MW: ***p = 0.0002; FT: ns, p = 1.0. **D)** *MW*: ***p = 0.0001; FT: ns, p = 0.7387. **E)** *MT:* ns, p = 0.7183; FT: ns, p= 1.0. **F)** MW: ns, p = 0.5361; FT: ns, p = 0.6417.

We next investigated if other perturbations of the pigmentation pathways could affect mosquito susceptibility to *P. berghei* infection. For this, we used a loss-of-function mutant of the hydrolase Tan involved in the homeostasis of the two pigment precursors dopamine and NBAD (**Fig. 1**). In contrast to *yellow(-)KI* mutants, *tan(-)KI* mosquitoes displayed similar parasite loads as wild type controls raised together upon infection with high doses of *P. berghei* gametocytes (medians: 70 vs 58, respectively, **Fig. 4F**).

### Melanization response of *yellow(-)KI* mosquitoes against *P. berghei* oocysts

Ookinetes that are killed upon traversal of the midgut epithelium are either lyzed or melanized, an immune response that is highly dependent on the mosquito genetic background (Volz *et al*., 2006). Melanization is triggered through the specific activation of prophenoloxidases (PPOs) in proximity of the parasite. In their activated form, PPOs oxidize monophenols and o-diphenols to o-quinones that are precursors of melanin (Nappi and Christensen, 2005). Since Yellow is believed to affect the synthesis of the quinone dopachrome, a precursor of eumelanin, we hypothetized that in the absence of Yellow, this molecule is absent or treduced in its concentration, potentially altering pathogen melanization. Since melanizing activity in *Ngousso* and *G3L* is very low, we introgressed the *yellow(-)KI* transgene into the *L3-5* line that is resistant to infection with *P. berghei* and displays a strong melanization response (Collins *et al*., 1986). Females carrying the *yellow(-)KI* transgene were backcrossed 6 times with L3-5 males (**Fig S2A**). Female *yellow(-)KI* carriers of the sixth generation (BC6) were crossed again with L3-5 males and allowed to lay eggs separately in single tubes to generate isofemale families. Each BC6 female was genotyped for *TEP1* (**Fig. S2B**), and the progenies of 6 females homozygous for *TEP1R* were kept and mixed to ensure that all BC7 individuals were homozygous for *TEP1R*, an immune gene allele that is critical for the resistance and melanization phenotypes typical of L3-5 mosquitoes (Zheng *et al*., 1997). BC7 mosquitoes were intercrossed to establish a *L3-5* colony with a floating *yellow(-)KI* allele. Subsequently wild type and *yellow(-)KI-L3-5* females derived from this colony were infected with *P. berghei* and the numbers of live oocysts (EGFP-positive) and melanized ookinetes (black spots in brightfield) were counted 7-8 days after infection. Susceptible mosquitoes from *Ngousso* wild type were infected in parallel as a control, to confirm all infections were efficient and produced high and similar numbers of oocysts. Homozygous *yellow(-)KI* mosquitoes were fully resistant to *P. berghei* infection, as wild type L3-5 mosquitoes, and 100% of them also carried melanized ookinetes, however significantly fewer than wild types (medians: 67 vs 125, respectively) (**Fig. 5A,B**). These results indicate that, in the absence of Yellow, a larger proportion of killed ookinetes is eliminated through lysis instead of melanization. Still, melanized parasites in *yellow(-)KI* and WT appeared identical (**Fig. 5C**), suggesting that mosquitoes are able to melanize pathogens in the absence of Yellow.

**Figure 5.**
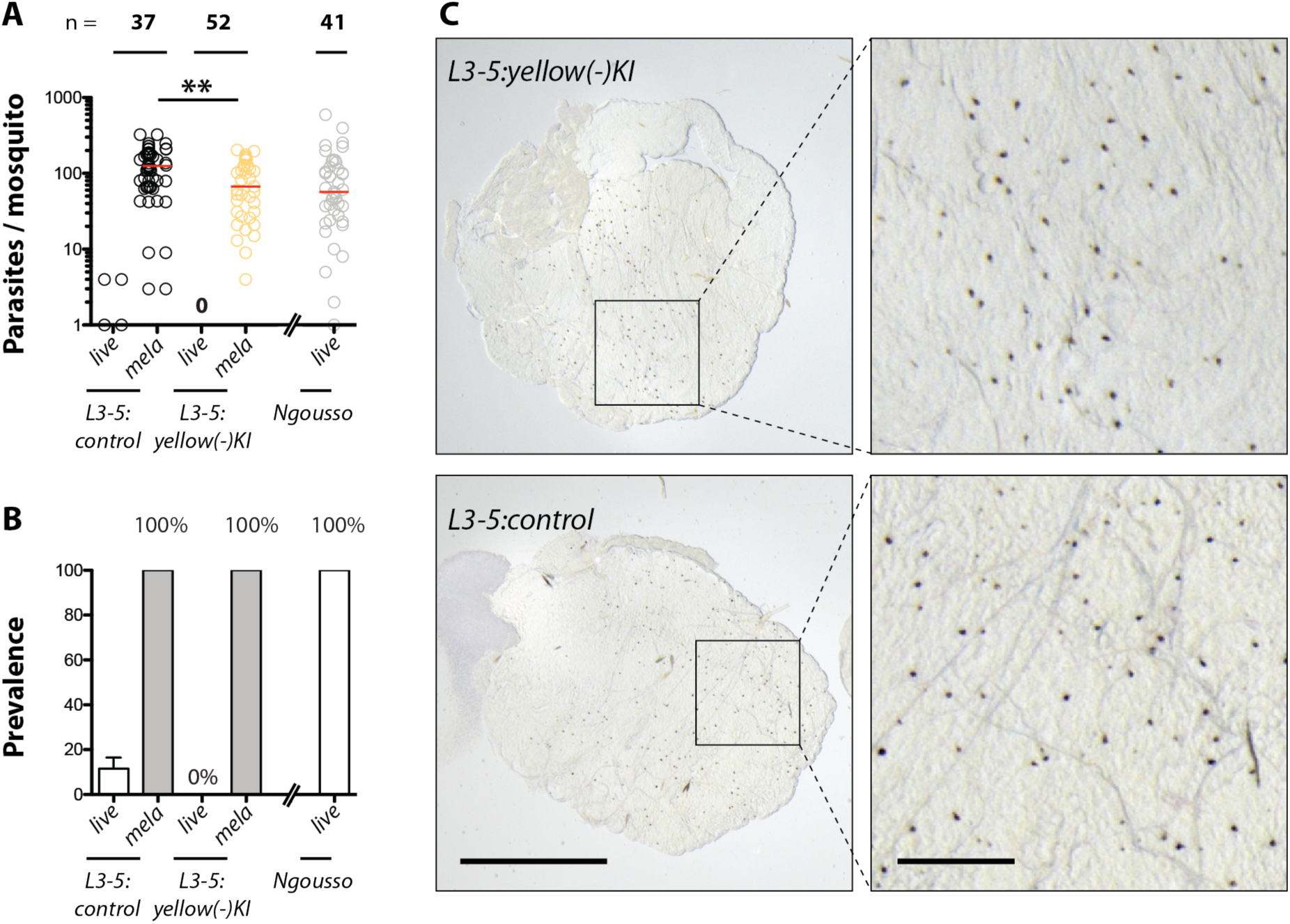
*yellow(-)KI* does not affect resistance to *P. berghei* but reduces ookinete melanisation in *L3-5* mosquitoes. **A)** Mosquitoes homozygous for *yellow(-)KI* and negative siblings of the resistant strain *L3-5*, and mosquitoes from the susceptible Ngousso strain were infected with fluorescent *P. berghei* (PbGOMO). Numbers of live (EGFP-positive) and melanized parasites at 7-8 days post infection are plotted, with red bars indicating medians. Two independent biological replicates. Statistical significance was tested using Mann Whitney test: **p = 0.0025. **B)** Percentage of mosquitoes carrying live and melanized parasites (prevalence) from the same infections as in (A). Shown is the mean and the standard error of the mean (SEM). **C)** Midgut of an infected L3-5 mosquito homozygous for *yellow(-)KI* in comparison to a transgene-negative sibling (control) seven days post infection. Melanized parasites appear as black dots. Scale bar: 500 µm (left) and 100 µm (right, zoom).

### Elevated parasite counts in *yellow(-)KI* females are independent of the complement factor TEP1

A major resistance factor to *Plasmodium spp.* in mosquitoes of the *A. gambiae* complex is the thioesther protein TEP1 (Blandin *et al*., 2004). TEP1 is a structural homolog of the human complement factor C3 and believed to act in a similar fashion, inducing lysis, phagocytosis or melanization through binding to foreign surfaces (Blandin, Marois and Levashina, 2008). To investigate whether increased *P. berghei* loads in *yellow(-)KI* mosquitoes could be due to a requirement of Yellow for TEP1 activity, we introgressed the *7b* transgene in the *yellow(-)KI* line. *7b* is an X-linked dominant transgene that triggers silencing of the endogenous *TEP1* locus (Pompon and Levashina, 2015). We reasoned that if the *yellow(-)KI* phenotype was dependent on TEP1, then *7b* and *yellow(-)KI;7b* mosquitoes would display similar susceptibilities to *P. berghei* infections, else *yellow(-)KI;7b* would be more infected. Virgin homozygous *7b* females and *yellow(-)KI* males were crossed and neonate F1 larvae were COPAS sorted for EGFP (marker of the *yellow(-)KI* transgene) and dsRed (marker of the *7b* transgene). Since both *7b* and *yellow* reside on the X chromosome, F1 mosquitoes carrying both transgenes were exclusively female. To select for recombinants harboring both transgenes on the same chromosome, F1 females were backcrossed to *yellow(-)KI* males and F2 offspring was again fluorescence sorted for dsRed and EGFP. Since male *Anopheles* mosquitoes possess only one X chromosome, F2 males positive for EGFP and DsRed had to be recombinants. These recombinant males were backcrossed to *yellow(-)KI* females and subsequent generations were fluorescence sorted for EGFP and DsRed until saturation. Of note, F1 males negative for *yellow(-)KI* but positive for 7b were backcrossed to 7b females to establish a separate colony with a similar genetic background as *yellow(-)KI;7b*. This colony served as control for all infection experiments with *yellow(-)KI-7b*. We first confirmed by Western Blot that TEP1 was depleted in *7b* and *yellow(-)KI;7b* mosquitoes (**Fig. 6A**). It appeared to be expressed at similar levels in wild type and *yellow(-)KI* mosquitoes, suggesting that *TEP1* expression and processing is not affected by Yellow depletion (**Fig. 6A**). Upon infection with *P. berghei*, *7b* mosquitoes carried higher parasite loads than wild type mosquitoes infected in different experiments but in similar conditions, as expected in the absence of TEP1 (medians: 239 vs 42, respectively, **Fig. 6B** and **4B**). Still, *yellow(-)KI-7b* mosquitoes were even more infected than *7b* mosquitoes (medians: 397 vs 239, respectively, **Fig. 6B**), indicating that the increase in susceptibility to *P. berghei* in *yellow(-)KI* mosquitoes is independent of TEP1 function. This also indicates that the high parasite loads observed in the absence of TEP1 do not reflect a maximal reachable infection level, since it can be further increased by removal of additional factors such as Yellow.

**Figure 6.**
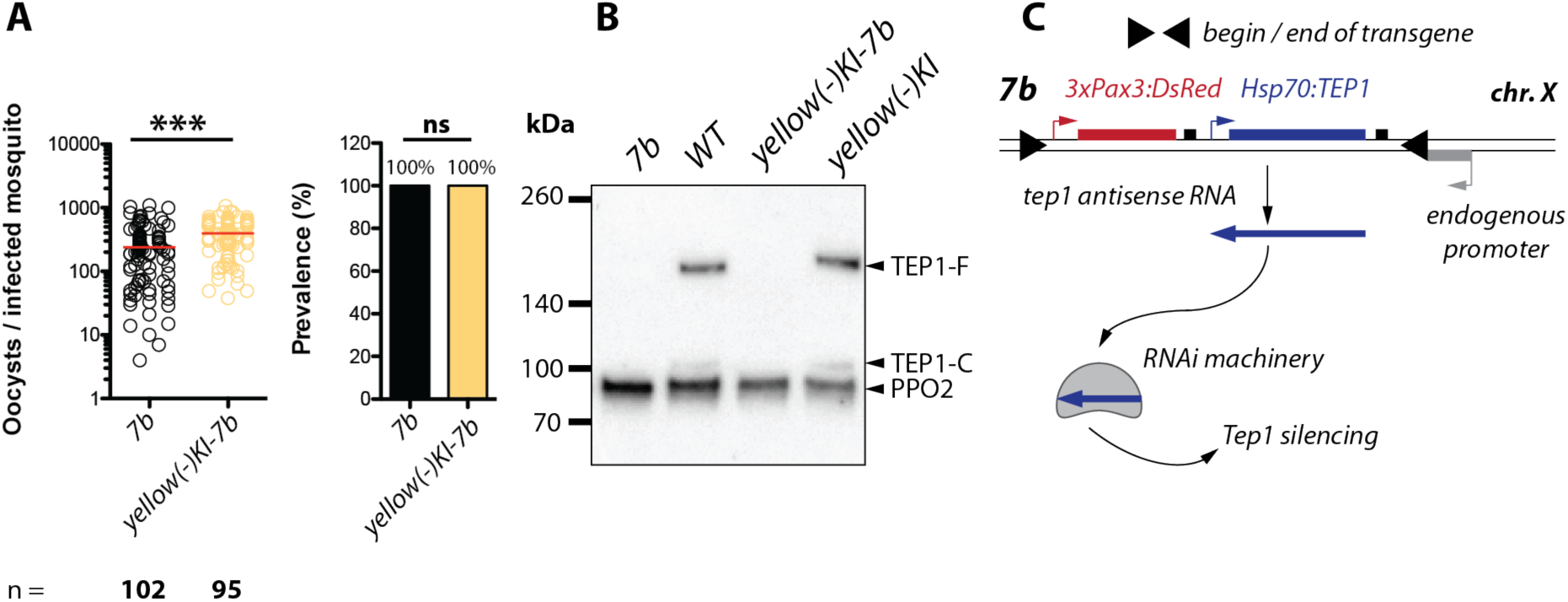
The antiparasitic effect of Yellow is independent of TEP1. **A)** *Yellow(-)KI* and *yellow(-)KI-7b* mosquitoes were infected with *P. berghei* (*PbGOMO*). Parasite load (oocysts per infected mosquito) with red lines indicating medians, and percentage of infected mosquitoes (prevalence, mean ± SEM) from three biological replicates. The number of analysed midguts is given below each genotype. Note the increased parasite burden in *7b* mosquitoes compared to controls expressing *TEP1* (see **Fig. 4**). Statistical significance was tested using Mann Whitney for parasite loads: ***p < 0.0001 and Fisher’s exact test for prevalences: not significant (ns), p = 1.0. **B)** Western blot on the hemolymph of *yellow(-)KI-7b* and *7b* mosquitoes probed against TEP1. Wild type and *yellow(-)KI* mosquitoes were used as positive controls expressing TEP1. An Anti-PPO2 (prophenoloxidase 2) antibody was used as loading control. **C)** Model of RNAi mediated knockdown of TEP1 in *7b* mosquitoes. A transgene composed of the *tep1* open reading frame under control of the *Hsp70* promoter from *D. melanogaster* causes a locus-specific effect leading to the transcription of a *tep1* antisense RNA subsequently inducing knockdown of TEP1 expression.

### Antibiotic treatment of *yellow(-)KI* mosquitoes rescues the infection phenotype

The increased parasite load in *yellow(-)KI* mosquitoes upon infection with *P. berghei* suggested a potential role of Yellow in the mosquito midgut where the first steps of parasite infection take place. Taking advantage of the fact that *egfp* is under control of the endogenous *yellow* promoter in the *yellow(-)KI* line, we examined whether *yellow* is expressed in this tissue. We did not detect EGFP fluorescence in *yellow(-)KI* mosquito midguts, including after blood feeding (**Fig. 7A**). This is coherent with our previous data showing that the *yellow* promoter activity is weak in adults and mostly restricted to the ovaries in females (Klug *et al*., 2022). Thus we hypothesized that the effect of Yellow on parasite development is indirect, either by influencing transcription and/or function of genes and/or proteins located downstream of Yellow, or through systemic metabolic changes induced by the absence of Yellow. Of note, the mosquito midgut microbiota composition influences the outcome of an infection and is regulated by a network of mosquito proteins for example FREP13 (fibrinogen-related protein 13) (Chauhan *et al*., 2020; Zaković *et al*., 2025). We thus tested whether the effect of Yellow absence on parasite development could be mediated by the microbiota. For this, *yellow(-)KI* and control mosquitoes were bred as a mixture and emerging adults were treated with different antibiotics supplied in 10% sucrose solution (**Fig. 7B**). Treatment of mosquitoes with an antibiotic cocktail (AB cocktail) composed of Penicillin, Amphotericin B, Streptomycin and Gentamicin completely rescued the infection phenotype in *yellow(-)KI* mosquitoes that displayed similar parasite loads as WT (medians: 122 vs 142, respectively, **Fig. 7C**). The same rescue was achieved upon treatment with streptomycin alone, an antibiotic acting on both Gram-negative and Gram-positive bacteria (median *yellow(-)KI* and WT: 78 and 57, respectively), but not when treated with Gentamicin alone, an antibiotic primarily acting on gram-negative bacteria (**Fig. 7C**). In this case, the phenotype was still present (median *yellow(-)KI* and WT: 167 and 85, respectively). Taken together, our results indicate that Yellow affects parasite development through its effect on the mosquito midgut microbiota, and that Yellow absence likely leads to the proliferation of Gram-positive bacteria that are beneficial, either directly or indirectly, to parasites.

**Figure 7.**
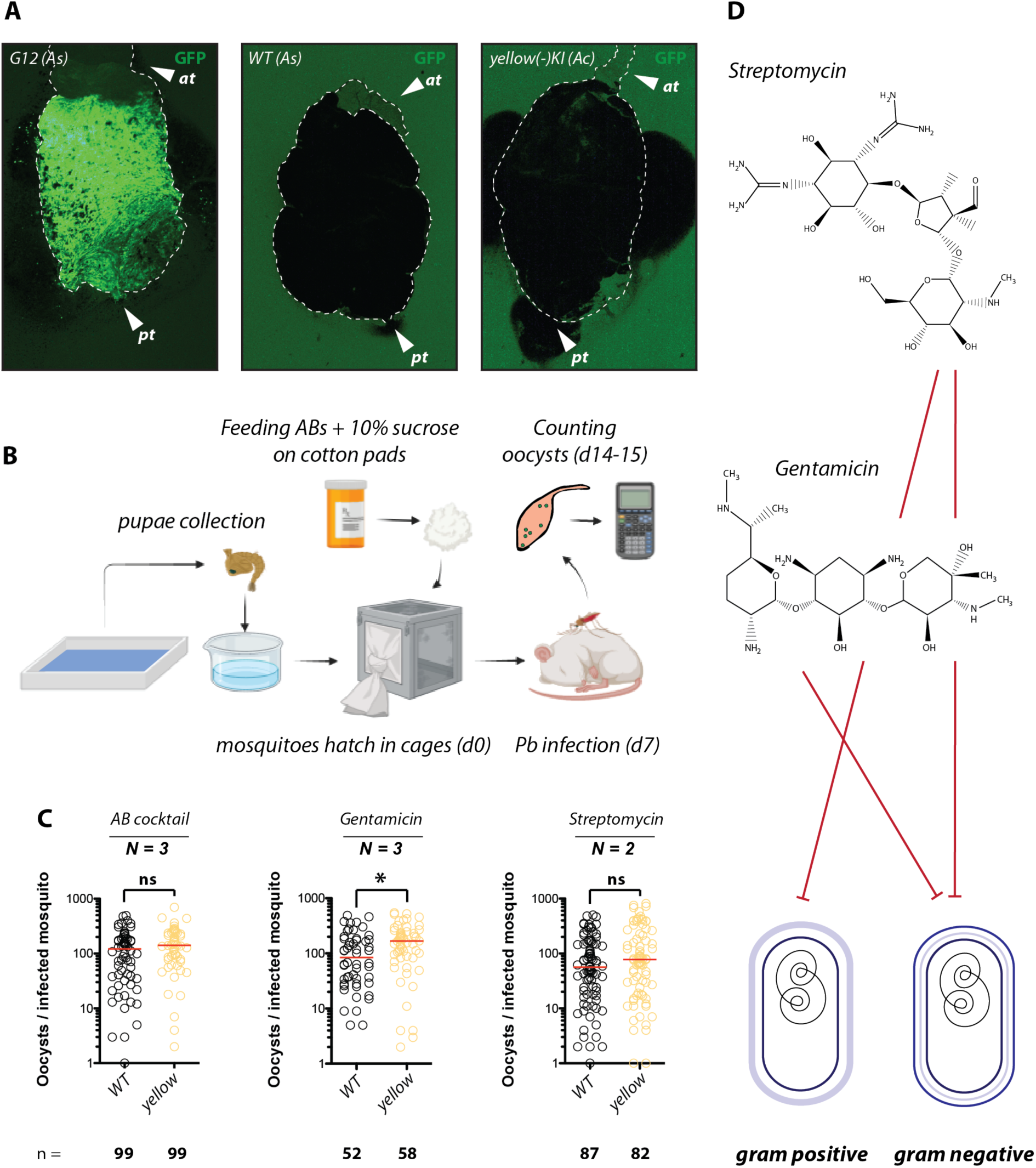
Differences in the oocyst burden between *yellow(-)KI* and control females disappear when treated with antibiotics. **A)** Midguts of *A. stephensi* (*As*) G12, *A. stephensi* (*As*) wild type (WT), and of *A. coluzzii* (*Ac*) *yellow(-)KI*, imaged to detect EGFP fluorescence 48 hours after blood feeding. The dashed white line highlights the shape of the midgut, while blood that is either in the intestinal lumen or has leaked into the surrounding area appears black. The anterior (at) or posterior (pt) end of the displayed guts are indicated within the images. G12 is a *A. stephensi* mosquito line expressing high levels of EGFP exclusively after blood feeding that was used here as positive control (Nolan *et al*., 2011). **B)** Juvenile stages of Y*ellow(-)KI* and wild type (WT) mosquitoes were bred together and allowed to emerge in the same cage. Adults were provided with cotton pads soaked with 10% sucrose solution supplemented with different antibiotics. They were infected with a high dose of *P. berghei* gametocytes seven to eight days after emergence. Oocysts were counted in the midguts on day 14-15 of life. The number of independent biological replicates (N) is given above each graph while the total number of dissected midguts (n) is given below. Statistical significance between *yellow(-)KI* and controls was tested using Mann Whitney (MW). The different antibiotics regimens are the following: **C)** streptomycin, penicillin, gentamicin and amphotericin B (AB cocktail). MW: not significant (ns), p = 0.1832; gentamicin. MW: *p = 0.0109; streptomycin. MW: ns, p = 0.2013. **D)** Activity profiles of gentamicin and streptomycin. Gentamicin targets predominantly Gram-negative bacteria while streptomycin targets both Gram-positive and Gram-negative bacteria.

## Discussion

Although *Yellow* was discovered more than a century ago, its precise role in pigmentation remains unclear. Current insights into *Yellow* function are largely based on visual observations and phylogenetic analyses, with no in-depth biochemical characterization. Moreover, phylogenetic studies have so far been restricted to insects, the only group in which this protein family is uniquely present. The only known related proteins, the major royal jelly proteins (MRJPs), constitute the primary protein source in honey bee royal jelly and have been implicated in the determination of queen development (Kamakura, 2011). While *Yellow* and MRJPs share some sequence similarity, available data to date suggest they perform distinct functions. We generated *yellow(–)KI*, a complete knockout allele of *yellow*, by inserting a reporter transgene into the first exon of the gene, and introgressed this allele into several genetic backgrounds to study its role in different mosquito life-history traits. In *A. coluzzii* (Ngousso background), the absence of *Yellow* significantly affected mosquito fitness, primarily through increased lethality and delayed development during juvenile stages. The resulting rapid elimination of the mutation in mixed populations may have been further exacerbated in our competition experiments with wild-type mosquitoes due to the experimental setup: most females—wild-type and mutant—likely mated with wild-type males, which emerged two days earlier than *yellow(–)KI* males and were thus ready to mate before their mutant counterparts. This would have accelerated the decline of the X-linked *yellow(–)KI* genotype in mixed cage populations. The increased lethality observed in some populations homozygous for the *yellow(–)KI* allele may stem from: (1) inbreeding effects due to the low number of founder transgenics, (2) susceptibility of specific genetic backgrounds to *Yellow* depletion, (3) increased susceptibility to bacterial proliferation in the culture water and (4) potential synergistic effects with other transgenes, notably *tan(–)KI* and 7b. Interestingly, in the *yellow(–)KI-aapp-DsRed* colony—a hybrid of Ngousso and G3L— although pupation was still delayed, both pupation rate and larval fitness were restored to wild type levels. These results suggest that the fitness impairment observed in competition experiments was largely due to delayed juvenile development in the absence of *yellow* expression, though additional effects on mating success, fertility and susceptibility to bacteria cannot be ruled out.

In the *yellow(–)KI* transgene, the *egfp* reporter is under the direct control of the artificial 3xP3 promoter, positioned immediately downstream of the endogenous *yellow* promoter. Consequently, its expression reflects not only classical 3xP3-driven patterns but also endogenous *yellow* activity. A similar pattern was reported for the *Sag(–)KI* allele, where *Sag(–)KI* and *Sag(–)EX* (post-3xP3 excision) showed comparable *egfp* expression pattern in the salivary gland, albeit expression was weaker in *Sag(–)EX* (Klug *et al*., 2022). This suggests that in this genomic context, the 3xP3 promoter amplifies *egfp* expression from the endogenous promoter, in addition to driving its expression in the eyes and ventral nerve cord. In *A. coluzzii*, we detected strong *egfp* signals in eggs, all larval stages, and pupae, whereas activity in adults was reduced and largely restricted to the ovaries in females (Klug *et al*., 2022). This differs markedly from *D. melanogaster*, in which *yellow* is predominantly expressed 46 hours after pupation, with minimal larval expression restricted to a subset of neurons and cells involved in mouthpart development. Consistent with this, *yellow* knockout in *D. melanogaster* has not been reported to affect larval development. Differences in *yellow* expression align with pigmentation patterns: while mosquito eggs and larvae are pigmented, fly eggs and larvae are mostly white and transparent. These pigmentation differences likely reflect adaptations to distinct breeding environments. The strong ovarian activity of the *yellow* promoter in mosquitoes correlates with the pronounced melanization of eggs after deposition. In various mosquito species, eggs with lighter cuticles exhibit lower hatching rates, indicating that pigmentation contributes to desiccation resistance (Farnesi *et al*., 2017; Noh *et al*., 2021). This is advantageous when oviposition occurs in small, ephemeral puddles prone to rapid drying. Similarly, the pigmented cuticle of mosquito larvae and pupae aligns with *yellow* activity and may serve as camouflage against predators. In contrast, fruit fly eggs are deposited in moist, decomposing material, where larvae quickly burrow, making both desiccation resistance and camouflage less critical. In adult *D. melanogaster*, *yellow* loss alters cuticle properties, and *yellow*-deficient males exhibit reduced sex comb stiffness, impairing mating success (Massey *et al*., 2019). Beyond visible phenotypes, *yellow(–)KI* mosquitoes displayed a consistent increase in susceptibility to *P. berghei*, with mutant mosquitoes carrying about twice as many oocysts as controls. This effect was independent of the complement factor TEP1 and the melanization response, as homozygous *yellow(–)KI-L3-5* females still melanized most killed ookinetes. Interestingly, knockout of *yellow* increased parasite burden in mosquito lines with the TEP1^S allele (linked to *Plasmodium* susceptibility) (**Fig. 4**) and in TEP1-depleted mosquitoes (**Fig. 6**), but *yellow(–*)*KI* mosquitoes expressing the TEP1R allele remained resistant (**Fig. 5**). Remarkably, the heightened *P. berghei* infection in *yellow(–)KI* mosquitoes could be rescued by antibiotic treatment—either with a cocktail or with streptomycin alone (effective against both Gram-negative and Gram-positive bacteria). Gentamicin alone (targeting mainly Gram-negative bacteria) reduced but did not eliminate the difference between mutants and controls. These findings suggest that *yellow(–)KI* alters gut microbiota composition or abundance, thereby influencing susceptibility to *P. berghei*. Possible explanations include reduced overall microbial load, similar to antibiotic treatment, or shifts in microbiota composition affecting *Plasmodium* colonization. For example, changes in gut pH and tryptophan catabolism mediated by *Asaia borogensis* and *Pseudomonas alcaligenes* can impact *Plasmodium* oocyst numbers (Wang *et al*., 2021; Feng *et al*., 2022), while different microbial communities correlate with opposite outcomes of *P. falciparum* infection (Zaković *et al*., 2025). Here, the lack of increased *P. falciparum* susceptibility suggests that microbiota changes resulting from *Yellow* depletion affect *P. berghei* and *P. falciparum* differently. Further research is needed to elucidate how a pigmentation gene like *Yellow* influences mosquito microbiota. The reduced parasite melanization observed in *yellow(–)L3-5* females suggests alterations in melanin precursor concentrations and/or composition. Overall, our results indicate that in *Anopheles*, loss of *Yellow* may have two major effects: (1) systemic changes in pigmentation precursors, and (2) altered gut microbiota density and/or composition, possibly as a downstream consequence of (1), since *yellow* is not expressed in the gut. This underscores the complex interplay between genetic traits, physiology, gut microbiota, and pathogen susceptibility, and opens new avenues for exploring the role of *Yellow* in mosquito biology.

**Table 1:**
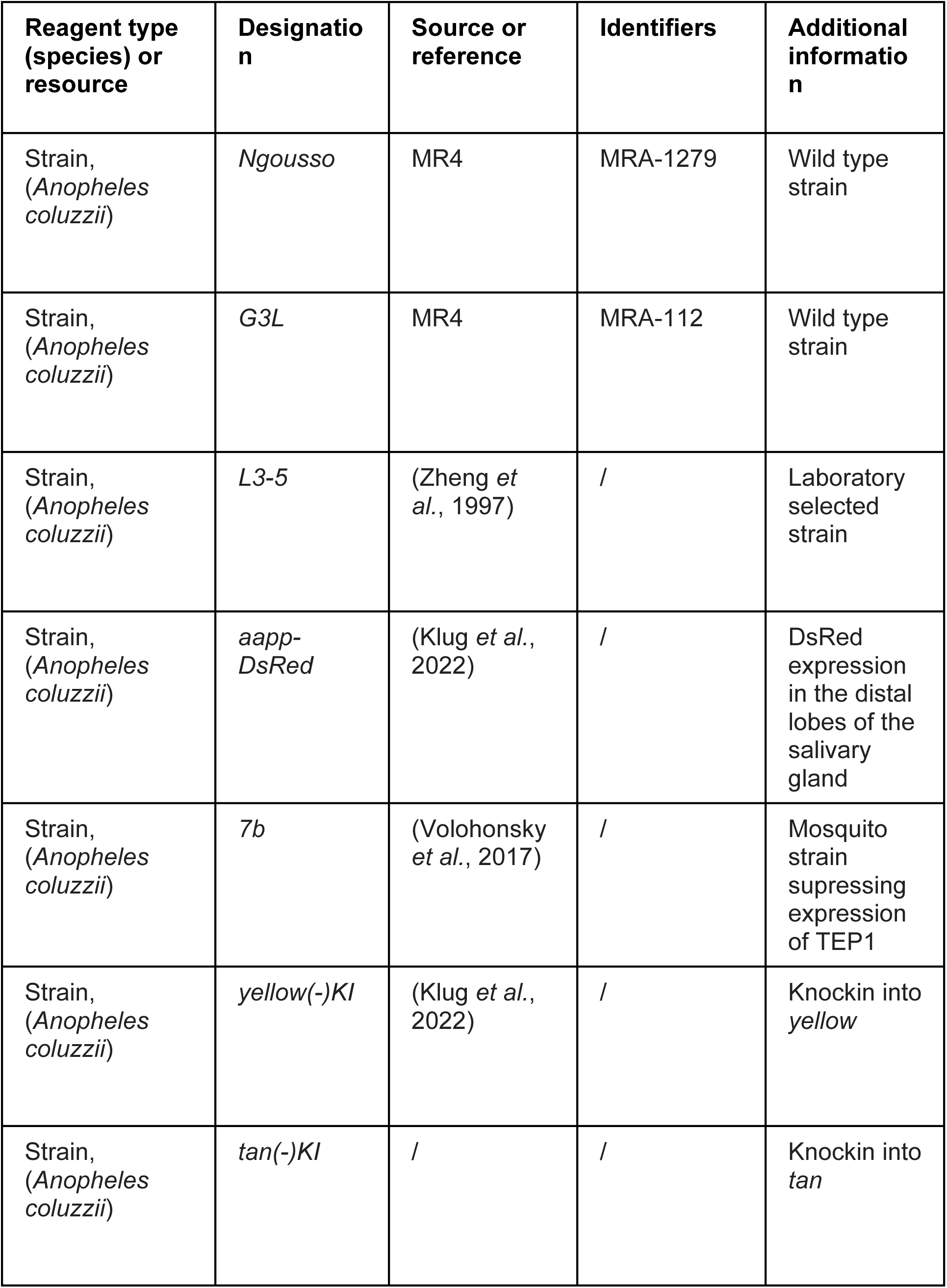
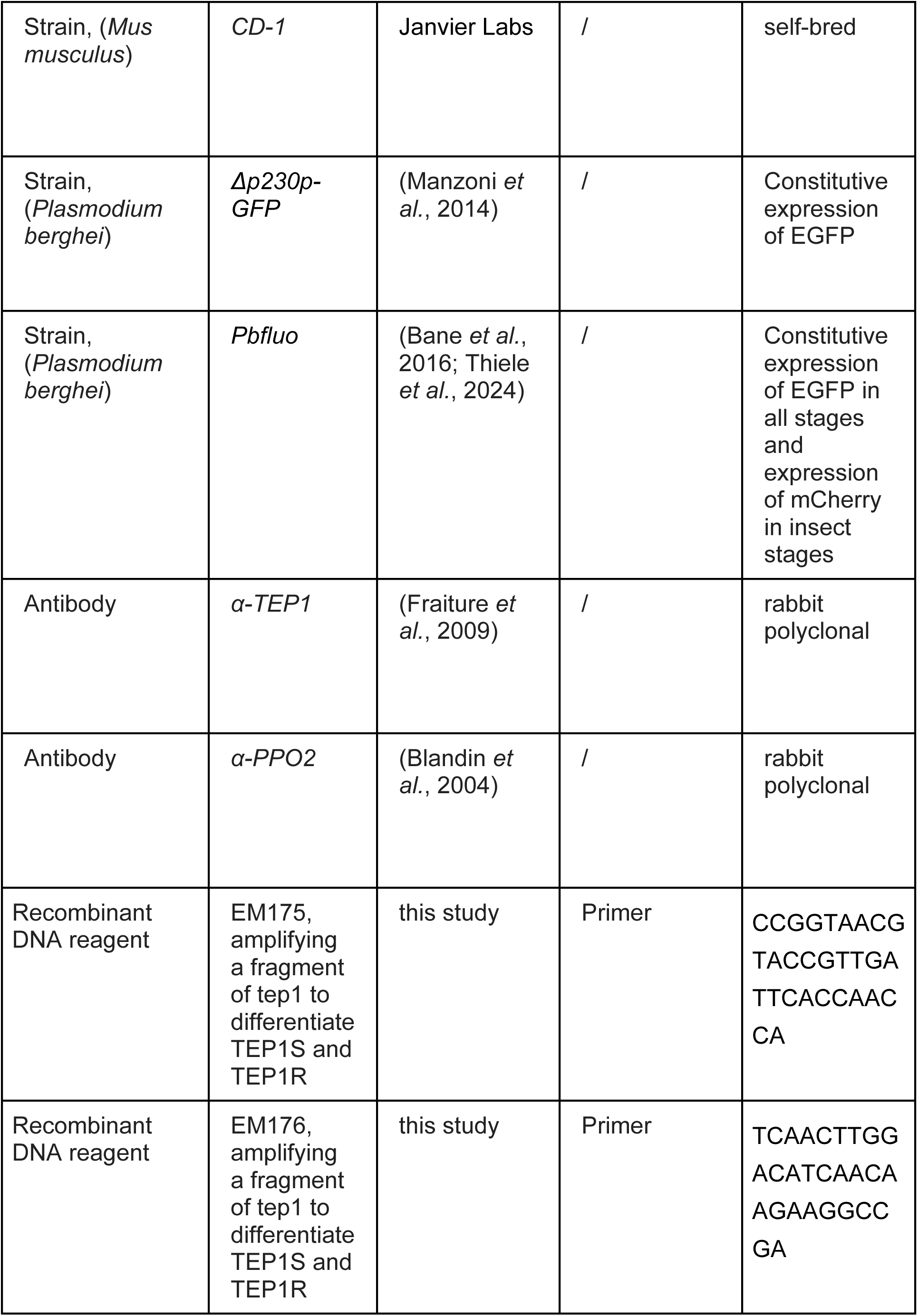
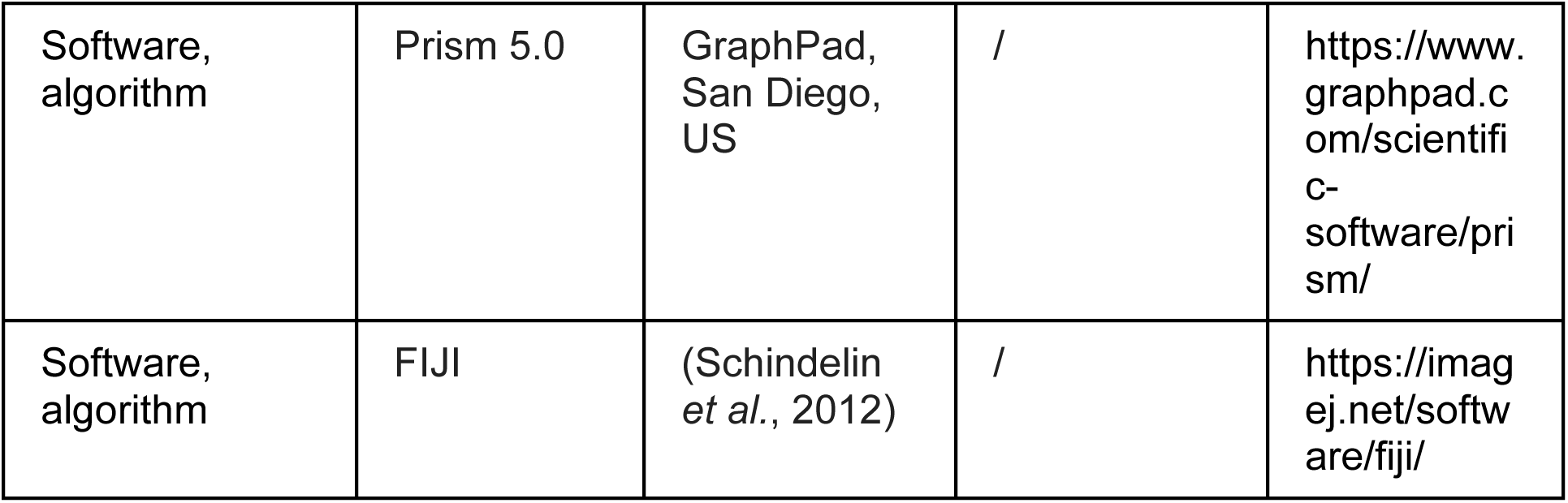
Key Resources.

## Materials and methods

### Breeding conditions

*A. coluzzii* (G3L, L3-5 and *Ngousso*) mosquitoes were kept at standard conditions (27-28°C, 75-80% humidity, 12-hr/12-hr light/dark cycle). Larvae were hatched in deionized water and fed with finely pulverized fish food (TetraMin). After pupation, pupae were collected in small glass dishes and transferred into netted cages. Hatched mosquitoes were fed with 10% sugar solution *ad libitum* using the capillarity of a cord, dipping in a tube filled with sugar solution. To propagate colonies, four to seven day old mosquitoes were blood fed for 10-15 minutes on anesthetized mice. Two days later mosquitoes were offered a glass dish with wet filter paper to allow egg laying. L1 larvae hatched after two days and were raised as described above.

### Mosquito transgenesis

The transgenesis of mosquitoes was performed following standard protocols (Volohonsky *et al*., 2015). Detailed information about the generation of *Yellow(-)KI*, *Tan(-)KI* and *aapp-DsRed* mosquitoes can be found in (Klug *et al*., 2022).

### Infection of mosquitoes with *P. berghei*

CD1 mice were infected through intraperitoneal injection of a frozen stock of *Plasmodium berghei* (*PbANKA*) parasites, PbGOMO (Manzoni *et al*., 2015), genetically engineered to constitutively express EGFP. Subsequently parasitemia was monitored by FACS (AccuriC6 SORP, Becton Dickinson). Once parasitemia reached 3-5 % mice were bled by cardiac puncture and infected blood passaged into naïve mice by intravenous injection into the tail vein. The number of injected mice depended on the number of mosquitoes planned to be infected while the number of transferred parasites depended on the planned infection date. Usually 2 x 10^7^ parasites were transferred into two mice to achieve a parasitemia of 2-3% at day three post injection. Four to seven day old female mosquitoes were transferred into paper cups (approximately 50 females per cup) the day before infection. On the day of the infection infected mice were anaesthetized and placed on the cup to allow mosquitoes to feed for approximatively 15 minutes covered by a dark blanket. Subsequently mosquitoes were anaesthetized on ice and non-fed females were removed. Infected mosquitoes were kept at 21°C, 60% humidity with a 12 hours light, 12 hours dark cycle. To achieve different infection levels mice which had been intravenously infected with *P. berghei* were either fed two days (low infection regime) or three days (high infection regime) after passage.

### Introgression of *yellow(-)KI* into *L3-5*

Virgin homozygous *yellow(-)KI* females were crossed with virgin L3-5 males to obtain heterozygous F1 *yellow(-)KI* mosquitoes inheriting 50% of the genetic background of each parental strain (*Ngousso* and L3-5). Virgin F1 females were crossed with L3-5 males to enrich for L3-5 specific genetic traits. Backcrossing was repeated for six generations while the *yellow(-)KI* transgene was selected by isolating only EGFP positive females through COPAS (Union Biometrica) sorting. In a last step of the introgression process, *TEP1R* homozygous females were isolated. This was necessary because this *TEP1* allele contributes strongly to the highly refractory phenotype of L3-5 mosquitoes against *Plasmodium spp.* infection. To create a mosquito colony expressing exclusively TEP1R, females from backcross six with unknown TEP1 genotype were mated with L3-5 males (which are homozygous for TEP1R) and blood fed. After two days, females were isolated in single *Drosophila* breeding tubes containing a piece of wet filter paper to allow egg laying. The offspring of selected females that laid eggs was raised as described above and the genomic DNA of founder females was extracted and used for TEP1 genotyping by PCR with primers EM175 / EM176. The generated PCR product (934 bp) was digested with the restriction enzyme *NcoI* (Thermo Fisher Scientific) for one hour at 37 °C, before gel electrophoresis. *NcoI* cuts specifically TEP1S but not TEP1R resulting in one fragment (undigested PCR product; 934 bp) if mosquitoes are homozygous for TEP1R, three fragments (digested PCR fragments: 783 and 151 bp as well as the full-length fragment of 934 bp) if mosquitoes are heterozygous for TEP1R and TEP1S and two fragments (completely digested PCR product; 783 and 151 bp) if mosquitoes are homozygous for TEP1S. The offspring of females homozygous for TEP1R was self-crossed to generate the homozygous *yellow(-)KI* line in the L3-5 background.

### Generation of *yellow(-)KI-7b* and *yellow(-)-aapp-DsRed* mosquitoes

Virgin 7b females and *yellow(-)KI* males were crossed and the F1 was COPAS sorted for EGFP (marker of the *yellow(-)KI* transgene) and dsRed (marker of the 7b transgene). Since the 7b transgene as well as *yellow* reside on the X chromosome, F1 mosquitoes heterozygous for both transgenes were exclusively female. To select for recombinants that harbour both transgenes on the same X chromosome, F1 females were backcrossed to *yellow(-)KI* males and the F2 offspring was fluorescence sorted for dsRed and EGFP. Since male *Anopheles* mosquitoes possess only one X chromosome, F2 males positive for EGFP and DsRed were recombinants possessing both transgenes on one X chromosome. Subsequently, recombinant males were backcrossed to *yellow(-)KI* females and following generations were fluorescence sorted for EGFP and DsRed until saturation. Of note, F1 males negative for EGFP (negative for *yellow(-)KI*) but positive for 7b were backcrossed to 7b females to establish a separate colony with a comparable genetic background as *yellow(-)KI-7b*. This colony served as control for all infection experiments in which *yellow(-)KI-7b* was used. To confirm absence of TEP1 protein, hemolymph of *yellow(-)KI-7b*, *yellow(-)KI* and control females was probed with αTEP1 antibodies on western blots which revealed complete absence of TEP1 in *yellow(-)KI-7b* females while *yellow(-)KI* females expressed TEP1 in levels comparable to the control (**Fig. 6B**). To generate *yellow(-)KI-aapp-DsRed* mosquitoes, virgin *yellow(-)KI* females were crossed to virgin male aapp-DsRed mosquitoes. The F1 generation of this cross was self-crossed and F2 larvae were sorted using the COPAS to keep only EGFP fluorescent larvae (marker for *yellow(-)KI*). EGFP positive larvae were treated with puromycin as described (Volohonsky *et al*., 2015). The selection process was repeated until EGFP reached saturation and lethality after puromycin treatment could no longer be observed. To exclude the presence of *yellow(-)KI* females heterozygous for *aapp-DsRed*, which would be sufficient to confer resistance to puromycin, 20 freshly mated and blood fed females were separated in single plastic vials for egg laying. The offspring of these females was raised and self-crossed to obtain the F2 generation without any selection. In case founder females were heterozygous for *aapp-DsRed*, *yellow(-)KI* females without *aapp-DsRed* must be present in the F2 generation. To exclude contamination with specimens lacking *aapp-DsRed*, living females of each colony were screened for red fluorescent salivary glands with a Nikon SMZ18 Stereomicroscope (Klug *et al*., 2022). Families with DsRed negative females were excluded from further breeding while colonies displaying only DsRed positive specimens were pooled to create a homozygous *yellow(-)KI-aapp-DsRed* line.

### Assessing differences in pupal development of *yellow(-)KI* and control mosquitoes

*Yellow(-)KI* and control (*Ngousso*) mosquitoes were reared in synchrony as two separate colonies. Newly hatched L1 larvae of both colonies were sorted using COPAS in batches of 50 larvae each. Mixed cultures of control and *yellow(-)KI* larvae were seeded in equal proportions, with 50 larvae for each genotype. Larvae were reared according to standard conditions. From day 7 to day 10, all emerging pupae were collected once a day and screened for the presence of EGFP using a Nikon SMZ18 Stereomicroscope with the respective fluorescence filter. This was necessary to clearly distinguish control (negative for EGFP) from *yellow(-)KI* pupae (positive for EGFP).

### Estimating fitness of *yellow(-)KI* lines

*Yellow(-)KI*, control (*Ngousso*), *yellow(-)KI-7b*, *yellow(-)-app-DsRed* and *yellow(-)KI;tan(-)KI* mosquitoes were reared in synchrony as separate homozygous colonies. Newly hatched L1 larvae of all colonies were sorted using COPAS in batches of 200 larvae each and seeded in separate pans. For each experiment (biological replicate) four pans (technical replicate) per genotype with 200 larvae each were reared. Larvae were grown in standard conditions. Between day 7 and day 10 emerging pupae, dead L4 larvae and dead pupae as well as living pupae with moulting defects were collected and counted. Experiments were terminated at day 11. The few remaining larvae were often small and appeared sick, so it was assumed that they would not continue to develop.

### Survival experiments

Four to seven day old homozygous *yellow(-)KI* and control females were separated in “survival” cups and kept at 21°C, 60% humidity and 12 hours light, 12 hours dark cycle. Survival cups were made from normal and conic drinking cups made from paper. The bottom of normal drinking cups was replaced with a glued conic cup to create a bottom with a pointed end. Subsequently the tip of the conic cup was removed and the resulting hole was closed with cotton. Dead mosquitoes gathered at the tip of each cone and were easily removed by removing the cotton while limiting escape risk. “Survival cups” were placed in small glass dishes to keep them stable. Mosquitoes received 10% sugar solution *ad libitum* provided as soaked cotton pads covered by petri dishes on top of each cup. Cotton pads were moistened every two to three days and renewed every seven to ten days. Dead mosquitoes were counted and removed every day.

### Treatment of mosquitoes with antibiotics

*Yellow(-)KI* and control (*Ngousso*) mosquitoes were reared synchronously as two separate colonies. Newly hatched F1 larvae of both genotypes were mixed to roughly 2/3 yellow(-)KI and 1/3 control larvae per pan. This was necessary to compensate for the higher mortality of the *yellow(-)KI* genotype and still obtain similar numbers of hatching *yellow(-)KI* and control females. Larvae were raised in standard conditions until pupation. Pupae were collected in small glass dishes and allowed to hatch in netted cages. From hatching onwards a 10% sucrose solution was provided supplemented either with an antibiotic (AB) cocktail (Gibco, Invitrogen Antibiotic-Antimycotic solution: 100 U/mL Penicillin, 100 µg/mL Streptomycin, 0,25 µg/mL Amphotericin-B; supplemented with 100 µg/mL Gentamicin (Sigma-Aldrich)) or exclusively with 100 µg/mL Streptomycin or 100 µg/mL Gentamicin. For treatment with Gentamicin and Streptomycin only, both chemicals (Sigma-Aldrich) were used at the indicated concentrations. The antibiotic treatment used is based on previous studies (Chabanol *et al*., 2020), but concentrations were slightly elevated as only adult animals were treated and not larval stages. Mosquitoes were infected 4-5 days after hatching as described before.

### Fluorescence imaging

Adults (**Fig. 2B**), eggs (**Fig. 2C**), larvae (**Fig. 2D, S2**), pupae (**Fig. 2D**) and midguts (**Fig. 5C**) were imaged using a Nikon SMZ18 Stereomicroscope with the respective filtersets for DsRed, EGFP and transmission light (no filter). The age and genotype of the specimens are indicated in the respective figures or in the figure legends, if applicable.

### RT-PCR

For RNA isolation 20 male and 20 female mosquitoes of wild type (*Ngousso*) and *tan(-)KI* were anaesthetized on ice and transferred to cryovials. 5-10 ceramic beads and 600 µl Trizol were added and homogenization of whole mosquitoes was performed using a Precellys (Bertin Technologies). Samples were either stored at −80°C or directly processed to isolate RNA. For RNA isolation 40 µl of p-Bromoanisol (BAN) was added. Tubes were vortexed for approximately 15 seconds and centrifuged for 20 min at 4°C and 16 rcf. After centrifugation the supernatant was transferred into a clean plastic reaction tube and 400 µl of isopropanol was added. Samples were mixed by gently inverting tubes several times before being centrifuged for 10 min at 4°C and 16 rcf. Subsequently the supernatant was discarded and pellets were washed with 500 µl of 70% ethanol. Samples were centrifuged for 5 min at 4°C and 16 rcf and the supernatant was discarded. Pellets were allowed to air dry for 5-10 min before being resuspended in 100 µl nuclease free water. Resuspended RNA was measured with a NanoDrop (Thermo Fisher Scientific) and 1 µg of RNA per sample was used for subsequent cDNA generation. DNAse digestion and reverse transcription was performed with the iScript gDNA Clear cDNA Synthesis Kit (Biorad, Hercules, California, United States) according to manufacturer’s protocols. The presence of TAN- (primer DK158 and DK159) and TEP1-specific cDNA (primer DK136 and DK137) was verified by perfoming PCRs using Phusion polymerase with 40 amplification cycles.

### Western blotting

Hemolymph was collected in denaturing protein loading buffer (Tris-HCl 0.35 M, SDS 10.3%, glycerol 36%, β-mercaptoethanol 5%, bromophenol blue 0.012%) by proboscis clipping from 10 mosquitoes ≥4 days after emergence. Hemolymph samples were denatured at 65°C for 5 min. Samples were separated by 7% SDS-PAGE. Protein membrane transfer, antibody incubations, and detection were carried out as previously described (Levashina *et al*., 2001).

### Ethics statement

The experiments complied with Directive 2010/63/EU of the European Parliament concerning the protection of animals used for scientific research. Our animal facility is approved under license number I-67-482-2 by the veterinary authorities of the Bas-Rhin department (Direction Départementale de la Protection des Populations). Authorization for animal use in this project, as well as for the production and use of transgenic organisms (bacteria, mosquitoes, parasites), was granted by the French Ministry of Higher Education, Research and Innovation under permit numbers APAFIS#20562-2019050313288887 v3 and 3243.

### Software and statistics

Statistical analysis was performed using GraphPad Prism 5.0 (GraphPad, San Diego, CA, USA). Data sets were either tested with a one-way ANOVA or a Student’s t test. A value of p<0.05 was considered significant. ChatGPT (GPT-5, OpenAI, San Francisco, CA, USA) was used to assist in refining the manuscript’s phrasing, spelling, and grammar.

## Acknowledgements

We thank Nathalie Schallon and Amandine Gautier for their assistance in mosquito rearing and *P. berghei* infections, and Ludvine Ramolu for technical support. We are also grateful to Lionel Brice Feufack Donfack for help with *P. falciparum* infections. We further thank the Mosquito Immune Responses (MIR) team for their support in mosquito breeding and for counting dead mosquitoes during survival assays.

---

**Figure S1.**
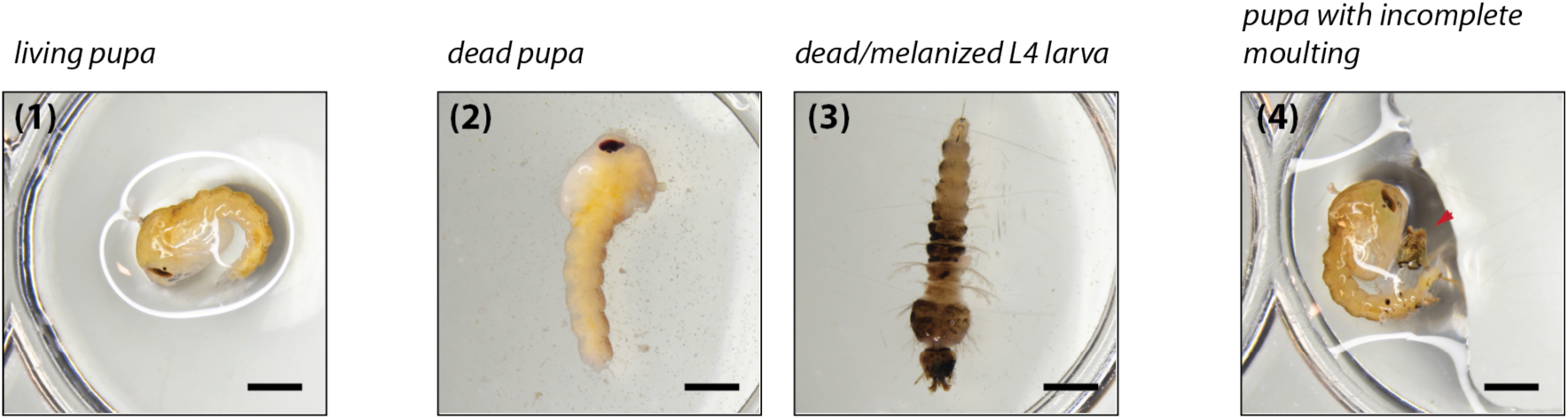
Lethality and moulting defects at larval and pupal stages in *yellow(-)KI* mosquito lines. Example images of (1) a living *yellow(-)KI* pupa, (2) a dead pupa with an unusual elongated posture, (3) a dead L4 larva - those often appeared black, likely due to melanization, and of (4) a living pupa with moulting defect. The last larval moult is still attached to the tip of the pupal abdomen (red arrowhead). Dead younger larvae are often difficult to detect as they are rapidly consumed by siblings. Scale bar: 2mm.

**Figure S2.**
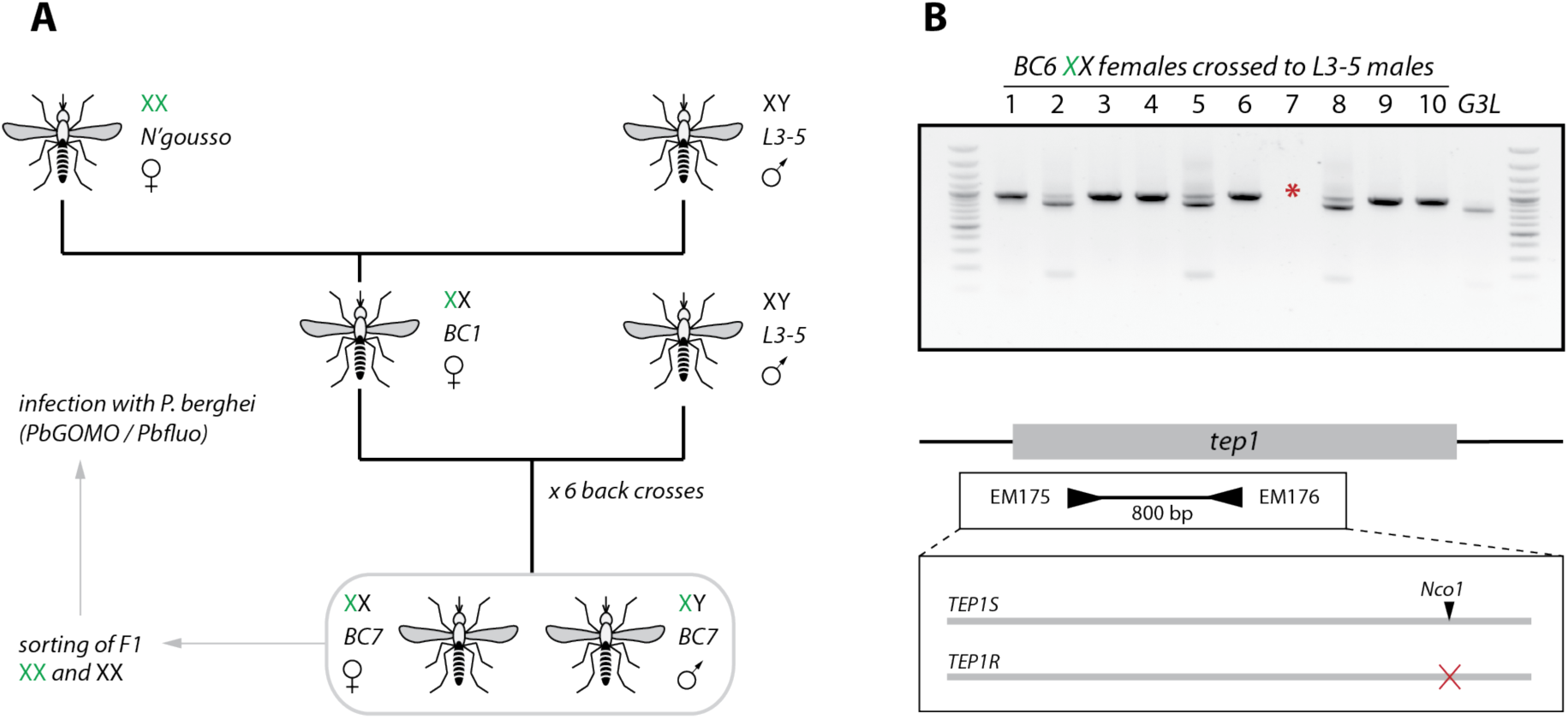
Introgression of the *yellow(-)KI* transgene into the *P. berghei* refractory mosquito strain *L3-5*. **A)** Crossing scheme to introgress the *yellow(-)KI* transgene into the *L3-5* genetic background. Females homozygous for *yellow(-)KI* were crossed with *L3-5* males. In the next generation, heterozygous BC1 *yellow(-)KI* females were sorted and crossed again with *L3-5* males. This crossing scheme was repeated for a total of seven times to yield a population of BC7 *yellow(-)KI* mosquitoes back crossed seven times to L3-5. **B)** During the last backcross, individual BC6 females were separated to lay eggs and genotyped in order to select progenies of homozygous *TEP1R* females only as founders of the *yellow(-)KI-L3-5* line. Indeed, *TEP1R* homozygosity is an essential feature of the *L3-5* phenotype. *TEP1* was genotyped using primers EM175 and EM176 and the PCR product was digested with *NcoI* cutting the PCR product of *TEP1S* once (two bands at 783 and 151 bp) while leaving PCR product of *TEP1R* uncut (one band at 934 bp). Females 1, 2, 4, 6, 9 and 10 were homozygous for *TEP1R* and their BC7 progeny was intercrossed. A G3L mosquito homozygous for *TEP1S* was used as control. For female seven (marked with a red asterisk) no PCR product was obtained.

